# Anterior thalamic nuclei neurons sustain memory

**DOI:** 10.1101/2021.08.25.457615

**Authors:** S. C. Barnett, L.C. Parr-Brownlie, B. A. L. Perry, C. K. Young, H. E. Wicky, S. M. Hughes, N. McNaughton, J. C. Dalrymple-Alford

## Abstract

A hippocampal-diencephalic-cortical network supports memory function. The anterior thalamic nuclei (ATN) form a key anatomical hub within this system. Consistent with this, injury to the mammillary body-ATN axis is associated with examples of clinical amnesia. However, there is only limited and indirect support that the output of ATN neurons actively enhances memory. Here, in rats, we first showed that mammillothalamic tract (MTT) lesions caused a persistent impairment in spatial working memory. MTT lesions also reduced rhythmic electrical activity across the memory system. Next, we introduced 8.5 Hz optogenetic theta-burst stimulation of the ATN glutamatergic neurons. The exogenously-triggered, regular pattern of stimulation produced an acute and substantial improvement of spatial working memory in rats with MTT lesions and enhanced rhythmic electrical activity. Neither behaviour nor rhythmic activity was affected by endogenous stimulation derived from the dorsal hippocampus. Analysis of immediate early gene activity, after the rats foraged for food in an open field, showed that exogenously-triggered ATN stimulation also increased Zif268 expression across memory-related structures. These findings provide clear evidence that increased ATN neuronal activity supports memory. They suggest that ATN-focused gene therapy may be feasible to counter clinical amnesia associated with dysfunction in the mammillary body-ATN axis.

**Highlights:**

- The mammillothalamic tract (MTT) supports neural activity in an extended memory system.
- Optogenetic activation of neurons in the anterior thalamus acutely improves memory after MTT lesions.
- Rescued memory associates with system-wide neuronal activation and enhanced EEG.
- Anterior thalamus actively sustains memory and is a feasible therapeutic target.

Optostimulation of anterior thalamus restores memory function after MTT lesions
Created with BioRender.com

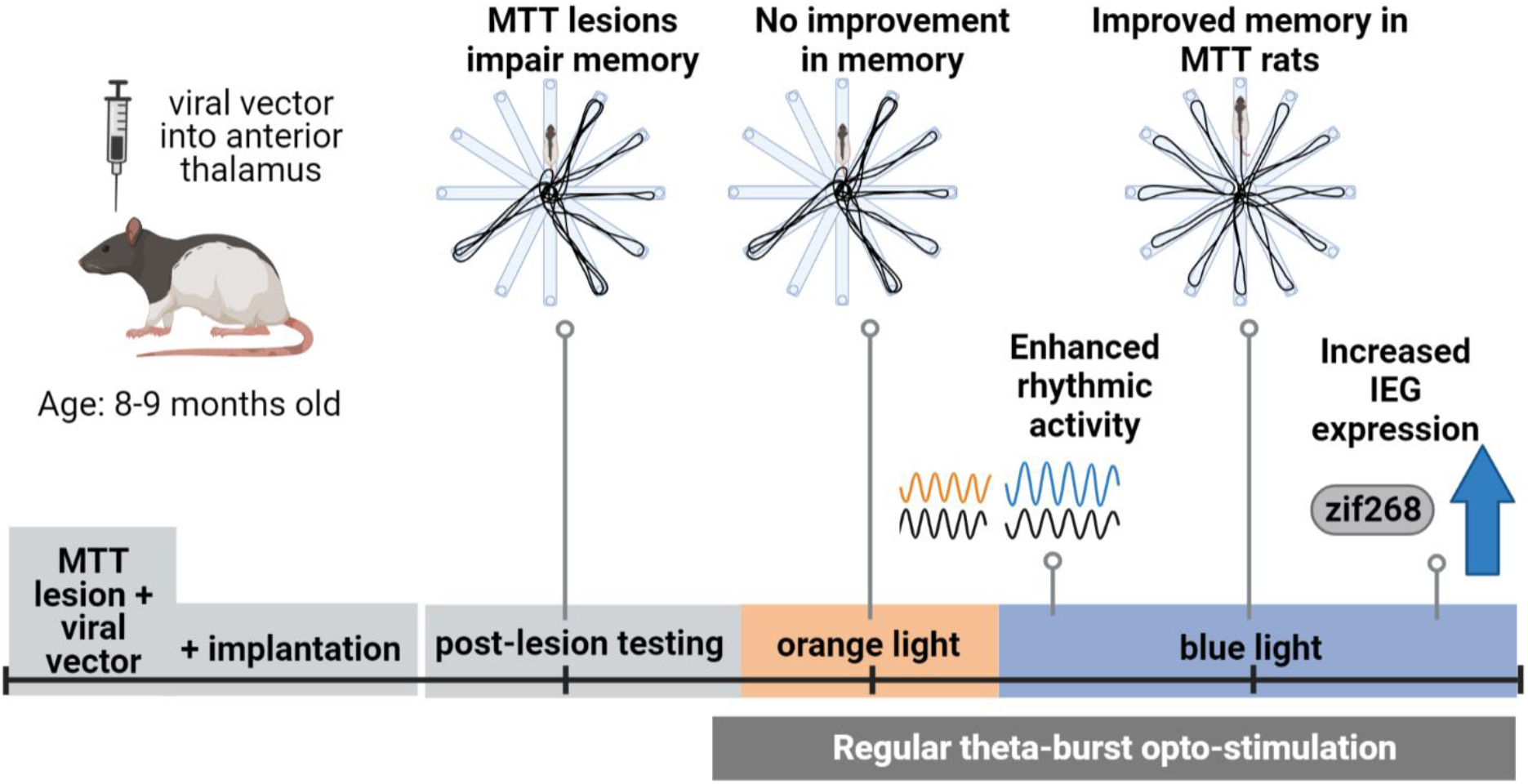

## 1. Introduction

Anatomical, clinical, and experimental lesion evidence suggests that the anterior thalamic nuclei (ATN) form a critical subcortical hub within a hippocampal-diencephalic-cortical network that supports memory function (Harding et al., 2000; Aggelton, 2014; Aggleton et al., 2016; Bubb, et al., 2017; Ferguson et al., 2019; Nelson et al., 2020; Nelson, 2021). Participation in a broad memory system is supported by many reports that ATN injury is also associated with metabolic and microstructural dysfunction in cortical and hippocampal structures (Reed et al., 2003; Caulo et al., 2005; Aggleton, 2008; Aggleton and Nelson, 2015; Dalrymple-Alford et al., 2015; Perry et al., 2018). Together, this evidence corroborates the idea that the ATN are responsible for more than the simple transfer of information to other key structures in the memory network (Wolff and Vann, 2019).

The negative impact of ATN lesions does not directly answer the question of whether ATN neurons actively facilitate memory, but there is some support for this claim from human studies. Functional magnetic resonance imaging data suggest that both ATN and mediodorsal thalamus activity is associated with memory in healthy adults (Pergola et al., 2013; Zotev et al., 2018). Memory performance in recovered Wernicke patients correlates with the degree of functional connectivity between the mammillary bodies (MB) and the ATN (Kim et al., 2009). This recovery might reflect information conveyed by the mammillothalamic tract (MTT), which constitutes axons from the MB that project to the ATN. More direct evidence comes from intracranial electrophysiological recordings made in epilepsy patients, which suggests that the ATN integrates information from diverse cortical sources during encoding to guide successful recall and that 50 Hz stimulation of the ATN improves working memory (Liu et al., 2021; Sweeney-Reed et al., 2021). In rats, however, there is mixed evidence for the impact of electrical stimulation of the ATN on memory. Hamani and colleagues reported that high current electrical stimulation at 130 Hz focused on the ATN impaired fear conditioning and nonmatching-to-sample in intact rats when stimulated during training; low current stimulation had no effect (Hamani et al., 2010). Low current stimulation improved delayed (four minute) visual nonmatching-to-sample performance in corticosterone-treated, but not saline-treated, intact rats when memory was tested at remote but not recent time points; this improvement was linked with increased hippocampal neurogenesis (Hamani et al., 2011).

Optogenetics allows for the selective stimulation of specific neuronal populations, whereas electrical stimulation activates all surrounding cells and fibers of passage. If ATN projections to memory structures actively support memory, stimulating these neurons optogenetically should impact the other structures within the distributed memory system. Moreover, we reasoned that selective stimulation could improve impaired memory caused by brain injury such as MTT lesions. MTT injury is the most consistent predictor of diencephalic amnesia due to thalamic infarcts (Carlesimo et al., 2011). In support of this clinical evidence, deficits in spatial working memory and temporal (i.e. recency) memory are found after experimental MTT lesions in rats (Aggleton, 2014; Nelson and Vann, 2017; Perry et al., 2018; Dillingham et al., 2015, 2021). The longevity of the deficit after MTT lesions in rats is uncertain, so we used a demanding 12-arm radial arm maze (RAM) and first confirmed that these lesions produce a long-lasting spatial working memory impairment. The majority of ATN efferents derive from glutamatergic neurons (Zakowski, 2017), so we next used a viral vector to transduce these specific neurons and render them responsive to light stimulation. Medial MB neurons, which project to the AV and the anteromedial subregion (AM) of the ATN, modulate their firing rate at theta frequency (Nelson et al., 2018). Similarly, there are theta-modulated cells in the AV and AM, which fire rhythmically with hippocampal theta rhythm (Jankowski et al., 2013; Zakowski et al., 2017). There is also evidence that the integrity of the MB-ATN axis influences electrophysiology across the extended hippocampal memory network (Dillingham et al., 2019, 2021). During spatial working memory testing, therefore, we used optogenetic theta-burst stimulation (TBS) in the dorsolateral region of the ATN, focused primarily on the AV, and we recorded electrophysiology to determine the theta-range power spectrum density (PSD) within, and coherence across, the ATN-hippocampal-prefrontal cortex axis (ATN-HPC-PFC axis). We compared the effects of optogenetic TBS of the ATN with both no stimulation and control stimulation by an ineffective light wavelength, while rats were tested in the RAM.

The most salient evidence that the MB-ATN axis impacts a network of memory-related brain regions is that MTT and ATN lesions reduce the expression of immediate early gene (IEG) activity in these distal brain structures (Aggleton, 2008; Dillingham et al., 2015; Loukavenko et al., 2016; Perry et al., 2018). We know that projections from the ATN to cortical regions and the hippocampal subiculum are almost exclusively ipsilateral in the rat (Mathiasen et al., 2017). So, we measured the IEG response in distal brain structures when rats received optogenetic stimulation of the ATN using effective (i.e. blue-light) TBS in one hemisphere and ineffective (i.e. orange-light) TBS simultaneously in the contralateral ATN. To avoid confounds related to the recruitment of these areas due to memory demands, we applied this stimulation while rats foraged for food in an open field, conducted 90 minutes before sacrifice, and removal of the brain to analyse Zif268 expression. Zif268 was selected as this marker is associated with spatial memory formation and long-term plasticity (Jones et al., 2001; Penke et al., 2014; Farina and Commins, 2016, Gallo et al., 2018) and has successfully revealed reduced IEG activity in the extended memory system following both ATN and MTT lesions (Dumont et al., 2012; Frizzati et al., 2016; Perry et al., 2018). Regions of interest for IEG analysis included key structures within the broader memory system that receive direct innervation from the ATN (Aggleton and Brown, 1999; Bubb et al., 2017; Nelson, 2021).

## 2. Methods

### 2.1. Animals

The experiment used 27 male Piebald Virol Glaxo cArc hooded rats, bred in the Animal Facility at the University of Canterbury. Rats weighed between 290-320 g and were aged 8-9 months at the time of surgery (Fig. 1). Twenty-three rats were randomly allocated to opsin-Sham or opsin-MTT lesion groups and 3 rats were assigned to a non-opsin MTT group. Rats were housed in mixed-condition groups of three or four rats per standard opaque plastic cage (50 cm length, 30 cm wide, and 23 cm high) in a vivarium. The vivarium lights were off between 8 am and 8 pm, when behavioural testing was conducted; observation of the rats confirmed that they remained relatively inactive during the lights-on period. Following surgery, rats were housed individually for seven to ten days. Food was available *ad libitum* just before surgery and during recovery. For behavioural testing, rats received restricted food access to maintain 85% of their free-feed body weight. Water was always available. All procedures were approved by the University of Canterbury Animal Ethics Committee, New Zealand (2016/24R) and comply with the ARRIVE guidelines.

**Fig. 1.**
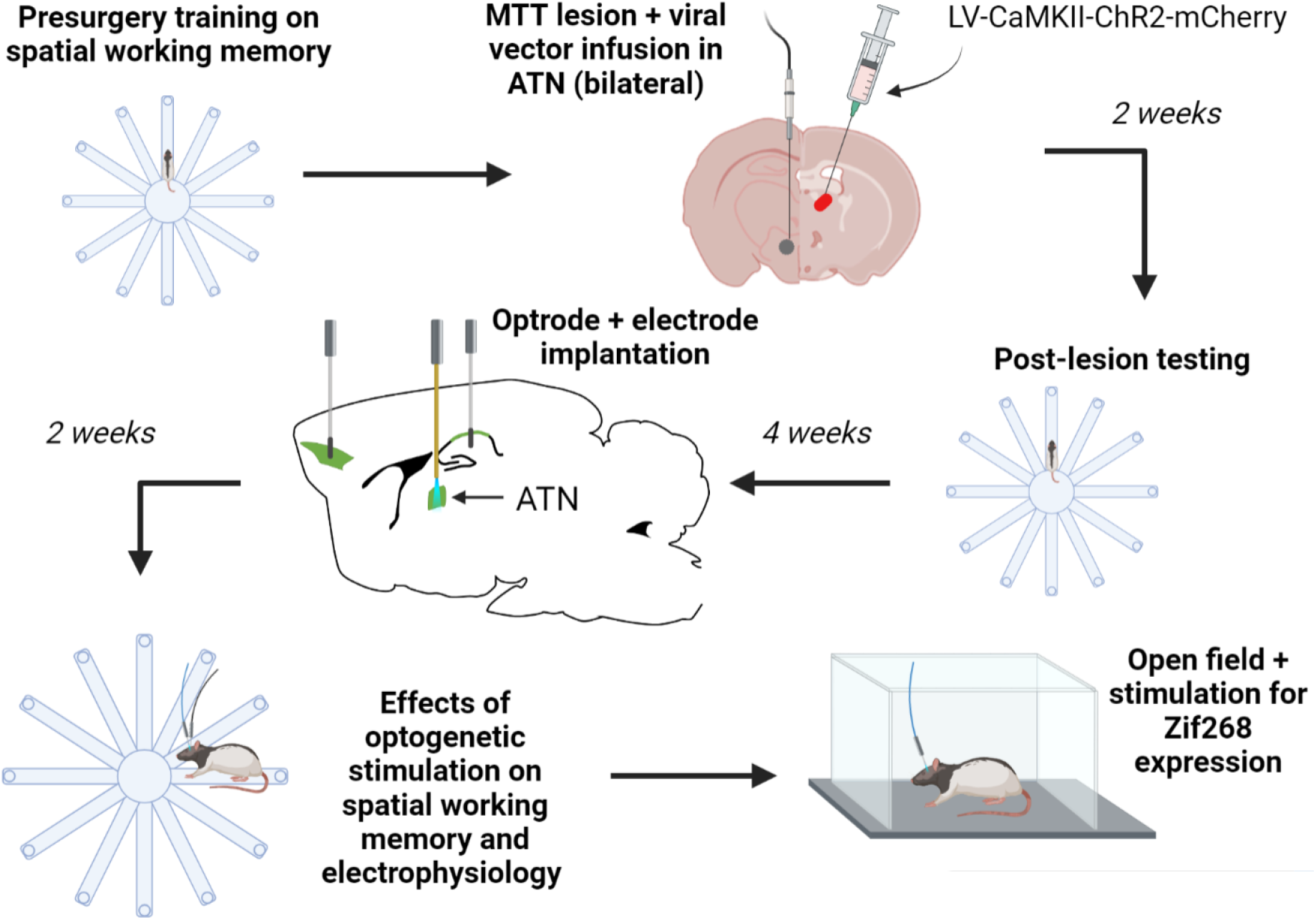
Temporal schematic of the key parts of the experiment. ATN = anterior thalamic nuclei; MTT = mammillothalamic tract. LV-CaMKII-ChR2-mCherry = lentiviral vector carrying channelrhodopsin construct; Zif268 = immediate early gene neuronal activity marker. Created with BioRender.com.

### 2.2. MTT lesion surgery

Rats were anesthetised with a ketamine (80 mg/kg i.p.) and Domitor (medetomidine, 0.35 mg/kg) mixture and placed in a stereotaxic apparatus with atraumatic ear bars (Kopf, Tujunga, CA) and dorsal skull surface held in the horizontal plane. On reaching anaesthesia, the rat was given Carprofen (s.c. 5mg/kg) for pain relief and 1 mL of Hartmann’s solution for hydration (sodium lactate 0.9%, i.p.). Methopt Forte eye drops were applied and a moist gauze was placed just above the eyes. The scalp was shaved and disinfected with 4% chlorhexidine gluconate before local analgesia (s.c. 0.2 mL of 2 mg/mL of Mepivacaine; additional applied after incision). Core temperature was maintained by insulating the rat’s body. One of four anterior–posterior (AP) coordinates from bregma was used for corresponding bregma to lambda (B–L) distances: −0.245 mm for B-L ≤ 0.64 mm; −0.250 mm for B-L 0.65-0.68 mm; −0.255 mm for B-L 0.69-0.72 mm; and −0.260 mm for B-L > 0.72 mm. An RFG4-A Radionics TCZ radio frequency (rf) electrode (0.25 mm diameter, 0.3 mm long exposed tip) was lowered vertically through a craniotomy, ±0.09 mm lateral to the midline and −0.71 mm below dura. The lesion (Fig. 2A) was created by slowly raising the temperature of the tissue surrounding the electrode tip to 65°C, maintained for 60 s, before slowly reducing the temperature to normal; this procedure promotes consistency in lesion size with rf lesions. Sham rats received the same procedure except the electrode was lowered to 1.0 mm above the lesion site and no rf current was applied.

**Fig. 2.**
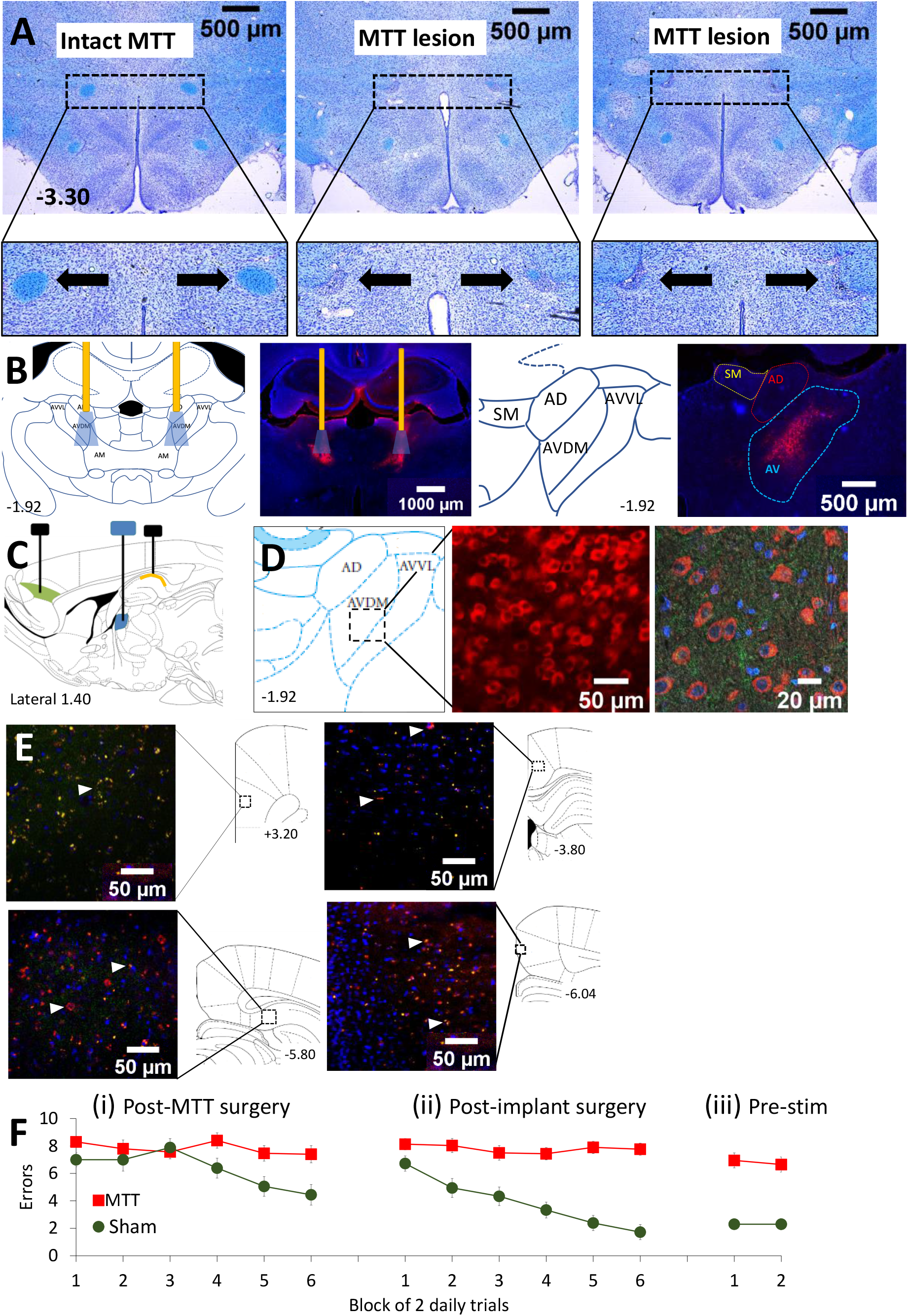
MTT lesions and viral transduction of ATN neurons. MTT lesions produced persistent spatial working memory deficits compared to Sham lesions. **A,** Photomicrographs of Luxol blue (myelin-specific stain) combined with cresyl violet (Nissl stain) showing the intact mammillothalamic tract (MTT) in a Sham lesion rat (left), one rat with 100% MTT lesion on the left and 83% lesion on the right (middle), and an example (right) from the 11 rats with 100% bilateral MTT lesions (two other rats failed the inclusion criterion of 80% bilateral damage). **B,** Placement of bilateral viral vector infusions of channelrhodopsin and optrodes focused on the AV subregion of the ATN. Red fluorescence indicates mCherry-positive transduced glutamatergic ATN neurons, which are predominantly transduced in the medial AV subregion and lateral AM subregion. Blue cones depict a conservative estimate of the distribution of effective light from the optic fibres (yellow bars), which can be up to 1.3 mm below the fibre tip. **C,** Sagittal schematic of electrode placements in dorsal HPC (CA1; yellow) and PFC (green), and optrode implant in the dorsolateral ATN (blue). **D,** Example of transduced neurons in the AV: the middle panel shows mCherry only, at 40x; the far-right panel at 60x shows co-expression of CaMKIIα immunofluorescence of glutamatergic neurons (green) and mCherry (red) based on co-expression (orange/yellow). Nuclei are stained with DAPI (blue). **E,** The mCherry and CaMKIIα co-expression (example arrow-heads) was confirmed in prefrontal (top left panel), subicular (bottom left panel), and retrosplenial (top and bottom-right panels) terminal regions. **F,** The 12 opsin-MTT rats showed persistent impairment in spatial working memory in the 12-arm maze compared to 9 opsin-Sham rats. Pre-stim refers to performance prior to any optogenetic theta burst stimulation. Abbreviations: AD, anterodorsal thalamic nucleus; AM, anteromedial subregion of the ATN; ATN, anterior thalamic nuclei; AV, anteroventral subregion of the ATN; AVDM, anteroventral thalamic nucleus, dorsomedial subdivision; AVVL, anteroventral thalamic nucleus, ventrolateral subdivision; HPC, hippocampus; PFC, prefrontal cortex. Error bars = ±SEM.

### 2.3. Viral vector surgery

About 40 mins after lesion surgery (i.e. same surgery session) rats received bilateral viral vector infusions targeted at the dorsolateral aspect of the ATN. The primary target was the AV subregion, but the spread of the viral vector and subsequent optic stimulation would be likely to include other ATN subregions. Opsin-MTT and opsin-Sham rats received two 0.24 μL infusions (i.e. 0.48 μL) per hemisphere of the lentivirus packaged with a channelrhodopsin construct (LV-CaMKII.hChR2(H134R).mCherry.WPRE; Optogenetics Resource Center, Stanford University), pseudotyped with vesicular stomatitis virus glycoprotein; the genomic viral titre was 5.34E+09 genomes/mL (from the Otago Viral Vector Facility, New Zealand). Lentiviral vectors were used because they have the dual advantage of limited spread from the target region coupled with the long-term expression of genetic constructs (Parr-Brownlie et al., 2015). Infusion coordinates were ±0.144 mm lateral; −0.525 mm and then −0.555 mm below dura. Infusions were made using a 10 μL nanofil syringe (World Precision Instruments, FL, USA) at a rate of 0.07 μL/ min. For infusions, the incisor bar was adjusted to −7.5 mm below the inter-aural line to minimise fimbria-fornix damage, which allowed the same AP coordinates as for MTT lesions just previously conducted with a flat skull. The syringe needle was lowered slowly to each coordinate (dorsal and then ventral on each side) and left *in situ* for five minutes for dispersion at each site. Three non-opsin MTT lesion rats (opsin control group) received LV-CaMKII.mCherry.WPRE, that is, missing the channelrhodopsin construct (6.26E+10 genomes/mL).

### 2.4. Electrode and optrode implant surgery

A second surgery, conducted 6 weeks later to allow ChR2 expression, was used to implant bilateral optrodes (optic fibre plus electrode) in the dorsal aspect of the ATN (Lateral, ±0.142 mm, DV −0.54 mm below dura). Bilateral dorsal hippocampal (HPC) electrodes (Lateral ±0.21 mm, DV −0.31 mm) and unilateral prefrontal cortex (PFC) electrodes (Lateral ±0.05 mm, DV −0.39 mm) were also implanted. The incisor bar was set at −7.5mm below the inter-aural line. The AP coordinates were −0.240 mm (AV), −0.420 mm (HPC) and +0.131 (PFC) for B-L <0.64 mm; an additional 0.005 mm was added for each corresponding B-L step listed in Section 2.2. Optrodes consisted of a pair of stepped stereotrodes glued with cyanoacrylate down the length of an 8 mm long optic fibre (200 μm diam, 0.66 NA; Doric Lenses, Quebec, Canada). The tip of the upper stereotrode extended ^~^100-200 μm below the tip of the optic fibre; the lower stereotrode extended a further ^~^200 μm (twisted 50 μm platinum-iridium wires, California Fine Wire, USA). Hippocampal electrodes consisted of four of these twisted wires at the same length and secured in place using the cyanoacrylate; the PFC stepped stereotrode consisted of pairs of the same twisted wire, but with one electrode placed 200 μm below the other. The twisted wires provided additional strength to the implant and the ability to select the channel showing minimal noise for electrophysiological recording. Ground electrodes were constructed from uninsulated 200 μm diameter pure silver wire (AM-systems). All implants were disinfected with 70% alcohol prior to implantation.

For implant surgery, anaesthesia was achieved using 4% isoflurane mixed with oxygen at a flow rate of 1500 mL/min, initially in an induction chamber. Once anaesthesia was reached, the rat was given a subcutaneous injection of Carprofen (5mg/kg) and 1 mL of Hartmann’s solution. The scalp was shaved, disinfected and local analgesia applied. Anaesthesia in the stereotaxic was maintained at ^~^2% isoflurane with an oxygen flow rate of 1000 mL/min. Five stainless steel anchoring screws were secured around the perimeter of the exposed skull. The sixth screw was placed in the orbital bone to act as the ground/reference electrode. The uninsulated silver ground wire was wrapped around the ground screw and then all remaining screws. Once all implants were secured, a head cap was constructed with successive applications of dental acrylic. The incision surrounding the head cap was cleaned with sterile saline and closed with sutures (if needed). Local anaesthetic cream was applied to the scalp area and the rat was given additional Hartmann’s solution. The rat was then administered 2 min of pure oxygen to facilitate recovery.

### 2.5. Electrophysiological recording and processing

When needed, rats were connected to the electrophysiology acquisition system (Open Ephys) plus the optogenetics stimulator via a head stage (Intan Technologies; CA) and extendable cables (PlexBright LED system, Plexon, Tx, USA) reaching a maximum length of 1.8 m were connected to a commutator that accommodated the attached Compact-magnetic optogenetic LED stimulators (OPT/Carousel Commutator 2LED-1DHST, Plexon, Tx). The commutator was positioned centrally above the hub of the 12-arm RAM. The rat’s headstage was plugged into the cable system just prior to testing and recordings were initiated once the rat had been placed in the maze hub. Electrode signals were referenced to the ground and recorded at 30 kHz for offline analysis of wideband local field potential activity (LFP). Recorded signals were down-sampled to 128 Hz, and notch (50 Hz) and bandpass filtered (1 −100 Hz).

Two 3 mm diameter infrared (IR) beams (each with emitter and receiver) were fixed just above the 3.0 cm walls of each of the 12 radial arms. One beam was located 15 cm along the arm to record the time of the rat’s entry; the second beam was located 12 cm from the end of the arm, just before the food well. The first beam break was used to reference epochs for analysis and to record arm entry. LFPs were extracted in 500 ms epochs for the rat’s last 8 correct choices on any given day. The use of correct arm visits facilitated the comparison between groups because rats with MTT lesions often did not achieve 12 correct arms whereas well-trained sham rats made relatively few errors. The 500 ms epoch was taken from 700 ms to 200 ms before the first beam break, to sample the initial act of making a choice in a previously unvisited arm. LFPs from ATN, HPC, and PFC electrodes were processed using NeuroExplorer version 5.128, Microsoft Excel, and SPSS. The LFP channel showing minimal noise or artefact per structure per rat was selected for all analyses. Filtered LFP channels were visually inspected and any data segments containing artefacts were removed before spectral and coherence estimates were calculated. Spectral analyses were performed separately for the ATN, HPC, and PFC in each rat in NeuroExplorer using a single taper spectral estimation (Hanning window) with 25% overlap. NeuroExplorer used a fast Fourier transform (FFT) for power and coherence calculations. FFTs were converted to power and normalised using a log transform (i.e., converted to log μV^2^ for analysis), extracted across a range of 2-14 Hz, and organised in Microsoft Excel before analysis in SPSS. For coherence analyses, the phase component was extracted from the FFTs, and coherence coefficients were calculated between recordings from electrode pairs for ATN-HPC, ATN-PFC, and HPC-PFC. Only correctly located electrodes were used to analyse power spectral density (PSD) within each structure, and only ipsilateral electrode pairs were used for coherence analyses.

### 2.6. Spatial memory and optogenetic stimulation

The 12-arm RAM used to test spatial working memory was located in the centre of a large, windowless room (4.0 m by 4.7 m) that provided ample spatial cues (door, tables, computer equipment, sink cabinet, and items on the walls). The maze was raised 70 cm above the floor and had 65 cm long, 10 cm wide aluminium arms with 3 cm high walls. Pulleys beneath the maze enabled the clear Perspex guillotine doors (10 cm by 28 cm) to be manually opened (by lowering them) or closed at the entrance to each arm around the perimeter of the 35 cm wide central hub. A clear Perspex barrier (19 cm high by 25 cm long) extended from the central hub along one side of each arm to prevent rats from jumping across arms. A wooden block (5 cm by 9 cm by 3 cm) was located at the end of each arm, with a recessed food well (3 cm by 1 cm) for 0.1 g chocolate food rewards; inaccessible chocolate under the wells provided constant olfactory cues.

Rats were habituated to the RAM and trained for 25 (±4) daily single-trial sessions prior to lesion surgery to achieve the criterion of a maximum of three errors per day for three consecutive days. The 12 arms were baited with one chocolate piece and the rat was placed in the central hub with all doors closed. After 5 - 10 s, all 12 doors were opened and the rats were allowed to select an arm (both hind legs completely inside an arm irrespective of reaching the food well). The rat was confined to the arm for ^~^10 s, irrespective of being correct or not, before being allowed back into the central hub where it was retained for 5 - 10 s before all of the doors were re-opened. Trials concluded when the rat had either visited all 12 arms or reached either 20 arm choices or 10 min had elapsed.

Following lesion surgery and brief re-habituation, testing was repeated for 12 consecutive days (Fig. 2, Fi). Because of the longer delay after the implant surgery, which permitted time for opsin expression, rats were re-habituated to the maze for three days before standard spatial working memory testing was assessed over another period of 12 days (Fig. 2, Fii), which included recording electrophysiology. The rat’s headstage was plugged into three cables, including the bilateral fibre optic patch cables, to familiarise them to the procedures used for later optogenetic stimulation.

Pre-stimulation RAM performance was again confirmed (Fig. 2, Fiii) prior to a 16-day period of optogenetic stimulation in the maze. First, we tested the effect of blue-light (465 nm, which activates the ChR2 opsins) using exogenously-triggered regular theta-burst 8.5 Hz stimulation (regular optogenetic TBS). This blue-light stimulation was compared with two control conditions. One control condition was orange-light regular TBS (620 nm, which does not activate ChR2). The second control condition was “no stimulation”, wherein rats were attached to all three cables but no optogenetic stimulation was delivered. The 16 days comprised 4 days of blue-light regular TBS; 4 days of orange-light regular TBS; and 8 days of no stimulation. The order of presentation for these three conditions was counterbalanced using the sequence [NOON][NBBN][NBBN][NOON], where N = no stimulation, B = blue-light TBS and O = orange-light TBS. This sequence enabled us to rule out order effects (first half vs second half and related interactions). Optogenetic TBS (for either blue or orange) was delivered via the PlexBright Optogenetic controller, triggered using parameters programmed into the Plexon Radiant software v2. The TBS used bursts of 3 x 5 ms pulses at 40 Hz, delivered at a frequency of 8.5 Hz, as a preliminary experiment found that this was the peak power spectral density for theta in the ATN and HPC for sham rats in the 12-arm RAM.

For optogenetic stimulation, the rat was connected to all three cables before being placed in the central hub of the maze. Once recordings appeared stable, optogenetic stimulation and/or electrophysiological recording and testing began. Stimulation was delivered continuously throughout the trial and terminated once all 12 arms had been visited or either 20 arm choices or 10 min had elapsed. At the beginning and end of every test day, the light output from the relevant LEDs (Plexon, Tx) was confirmed from the tip of the attached 0.50 NA patch cables (Power meter kit; Thorlabs). The power output at the optic cables using the blue LEDs was between 17.5-18.5 mW; the power output using the orange LEDs was 9-10 mW. These blue and orange outputs produce approximately similar variations in power density (irradiance mW/mm2) beyond ^~^150 μm in brain tissue and are estimated to influence neurons up to a distance of ^~^1.0 – 1.3 mm (https://web.stanford.edu/group/dlab/cgi-bin/graph/chart.php).

Five days later, we replicated the effects of the exogenously-triggered blue-light regular TBS. This used the same spatial working memory procedure and identical stimulation parameters. Rats were run for 4 days (two with blue-light regular TBS and two with orange-light regular TBS), using an OBBO counterbalanced design.

Starting the next day, we then assessed the effects of endogenously-triggered TBS in the same working memory task. For this new condition, the rat’s own hippocampal electrophysiological recording was filtered online (i.e., during testing) between 8-9 Hz by Open Ephys to yield an average of 8.5 Hz signal. The Open Ephys Phase Detector module triggered stimulation when the filtered HPC signal arrived at the upward zero-crossing or “rising phase” of the endogenous oscillation. This triggered the PlexBright Optogenetic Controller, via a Pulse Pal, to deliver a burst of 3 x 5 ms pulses at 40 Hz. This pattern of endogenously-triggered stimulation was regarded as “irregular” by comparison to the exogenously-triggered regular 8.5 Hz stimulation used earlier. The use of closed-loop methods for endogenous stimulation precluded us from recording the exact timings at which optogenetic stimulation was delivered. Rats were run in this way for 4 days, using an OBBO counterbalanced design.

### 2.7. Immediate early gene activation (Zif268) and Histology

At the end of the experiment, rats were habituated to being placed alone in a holding cage with regular bedding in a novel, quiet dark room for 90 mins per day for 3 days. On the next day, each rat was placed individually in a 50 cm high x 1 m x 1 m black open field with 20 chocolate pieces scattered across its base. Rats were connected to all three cables and allowed to explore for 6 minutes. During the last 5 minutes of this period, they were given exogenously-triggered blue light TBS in the ATN of one hemisphere and exogenously-triggered orange light TBS simultaneously in the contralateral side (counterbalanced across rats). After this, the rat was placed in the individual holding cage in the dark room for 90 min prior to anaesthetic overdose (sodium pentobarbital, 125 mg/kg) and perfusion. Once deeply anesthetised (cessation of plantar and tail pinch reflexes), the tip of each electrode was marked by passing 20 μA through the electrode for 20 s, to make a small lesion to identify electrode placement (including electrodes below the optic fibres), before transcardial perfusion with ^~^200 mL of chilled saline followed by ^~^200 mL of 4% paraformaldehyde in a 0.1 M phosphate buffer (PB) solution (pH 7.4). The head-cap and electrodes were carefully removed vertically to minimise brain damage and the extracted brain post-fixed in 4% paraformaldehyde solution overnight at 4 °C. Brains were transferred to a long-term storage solution (20% glycerol, 0.1 M PB) for a minimum of 48 h.

Coronal 40 μm sections were taken from approximately +3.70 mm to −9.16 mm from bregma using a sliding microtome with a freezing stage (ThermoFisher, Melbourne, Australia). Sections were collected in vials containing cryoprotectant solution (40% 0.1 M PB, 30% glycerol and 30% ethylene glycol) for long-term storage at −20°C. Four series were collected, with each series consisting of 4-6 sections. The first series of sections spanned from +3.70 mm to +1.20 mm from bregma, the second from −1.30 mm to −2.12 mm from bregma, the third from −2.12 mm to −3.80 mm from bregma and the fourth from −3.80 mm to −6.72 mm from bregma. For MTT lesion verification, one of six vials from the third series of brain sections was mounted onto gelatin-coated slides. Electrode and optrode verification used one vial from each of the first, second and third series of sections to cover the PFC, ATN, and HPC regions. Visualisation of LV.CaMKII.hChR2(H134R) mCherry.WPRE in the ATN used one vial from the second series of sections. The Zif268 processing used sections taken from one of each of the four vials.

### 2.8. Visualisation of MTT lesions and electrode/optrode placements

Luxol blue, a myelin-specific stain, was used to assess the integrity of the MTT pathways, with sections counterstained with cresyl violet (Nissl stain) to evaluate surrounding structures using standard procedures as used previously (Perry et al., 2018). Lesions were photographed under bright field at x10 on a Leica DM6 B upright microscope and DFC7000T camera (Leica Microsystems, Germany). The average cross-sectional area of the MTT at different AP levels in Sham rats was based on Luxol blue staining and estimated using the open-source Fiji software automated area calculation tool (ImageJ software, https://imagej.net/Fiji/Downloads). We used the Luxol blue staining to identify any intact MTT myelinated fibre area in the lesion rats to calculate sparing relative to the average intact MTT in the Sham rats; this sparing is less evident with only cresyl violet.

On removal from the cryoprotectant solution, sections were washed in PBS (3 x 10 min) and 0.1 M PBS-0.2% Triton (Tx), and incubated for 1 h in 0.1 M PBS-Tx 0.2% containing 10% normal goat serum (1:100 ThermoFisher, Melbourne, Australia) to block non-specific binding. Sections were transferred to well-plates for exposure to two primary antibodies (RFP antibody 5F8 rat for mCherry, RRID: AB_2336064, Chromotek, 1:1000; and mouse-monoclonal anti-CaMKIIα, RRID: AB_309787, Merck Millipore, 1:300) which were added to 0.1 M PBS-Tx 0.2% containing 5% normal goat serum. Sections were incubated in the primary antibody solution for 24 h at 4°C with gentle agitation. After this, sections were washed in PBS-Tx (2 x 20 min) and incubated for 4 hours at room temperature in the secondary antibody solution in the dark. The secondary antibody solution consisted of goat anti-rat Alexa Fluor-594 (Thermofisher, 1:1500, cat # A-11007) and goat anti-mouse Alexa Fluor-488 (Thermofisher, 1:1500, cat # A32723) added to 0.1 M PBS-Tx 0.2% containing 5% normal goat serum. The sections were then washed in PBS (2 x 10 min) and PB (1 x 10 min) before mounting on gelatin-coated slides in the dark. They were dried for ^~^30 min and coverslipped with Vectashield hardset mounting medium with DAPI (Vectorlabs, cat # H-1500-10). When dry, the slides were sealed with nail polish and stored at 4°C.

### 2.9. Zif268 immunohistochemistry

Sections were washed in 0.1 M PBS containing 0.2% Triton X-100 (PBS-Tx; 3 x 10 min). Endogenous peroxidase was blocked for 30 mins with a buffer solution (0.333ml of 30% hydrogen peroxidase, 5ml of 100% methanol, 1 ml PBS-Tx at 2%, and 3.667 ml distilled water). Sections were washed in PBS-Tx (3 x 10 min) before 72 h incubation at 4°C with rabbit polyclonal zif268 antibody (also known as Egr-1; 1:1500; Santa Cruz Bio, cat# SC-189) in PBS-Tx with 1% normal goat serum (NGS). After PBS-Tx rinses (3 x 10 min), sections were incubated in a biotinylated goat anti-rabbit secondary antibody (1:1000; Vector, cat# BA-1000-1.5) with 1% NGS in PBS-Tx for 24 h. They were then washed in PBS-Tx (3 x 10 min), before incubation in Extravadin (Peroxidase Conjugated; 1:1000; Sigma) in PBS-Tx with 1% NGS. Sections were rinsed in PBS (3 x 10 min), PB (3 x 10 min), washed in 0.05 M Tris buffer (pH 7.4; 2 x 5 min), and developed for 20 min in 0.05% diaminobenzidine (Sigma, Castle Hill, Aus) in 0.05 M Tris buffer with 30% hydrogen peroxidase added just prior to incubation. Washes in Tris buffer (3 x 10 min) were used to stop the reaction, followed by PB (3 x 10 min) and mounting on gelatinised slides. After drying, slides were put through graded alcohols (70%, 95%, and 100%) before clearance in xylene and cover-slipped with DPX mounting medium.

Regions of interest were captured with a 10x objective and automated cell counts made using the open-source Fiji software. To achieve semi-quantitative counts, the images were gray-scaled, the background-subtracted using rolling 20 light, the auto-threshold set using Max Entropy (built-in thresholding algorithm), and the image converted to binary. The same threshold was used across all rats. Labelled cell bodies were counted above the threshold with a circularity between 0.7 – 1.0 (circle, Circularity = 1). Cell counts were made blind to group conditions but were not stereological, providing a relative number of cells rather than absolute levels. Between four and six sections per hemisphere were analysed for each brain region. The cell count of Zif268-positive neurons in each sub-regional area was expressed as the number of cells per mm2, by dividing the total cell count by the corresponding area measured per um2, multiplied by 1,000,000.

### 2.10. Data analysis

Alpha was set at 0.05 for all statistical tests. Repeated measures ANOVAs were used for all behavioural analyses. Spatial memory performance after lesions (Sham versus MTT lesion), both prior to and after implant surgery, and the effects of optogenetic stimulation conditions, were analysed with ANOVA by extracting the linear and quadratic trends across successive two-day blocks. For Zif268 the distributions of raw cell counts were skewed (as often occurs with count data) and a square root transform was applied to achieve normal error distributions and so make the data suitable for ANOVA. Lesion and stimulation effects for these counts were analysed using ANOVA for each brain region separately, with additional repeated measures (subregions) as required. For analysis of electrophysiological data (using the last 8 correct choices per session, i.e., per single trial, for the reasons given above), there was the additional factor of electrode location (PSD) or electrode pair (coherence). There were similar running speeds for opsin-MTT and opsin-Sham rats, based on the time taken to run between the two beam breaks in the maze arms during correct arm choices in the first period of optogenetic stimulation (Group, F = 0.15, df = 1/20, P > 0.20); this suggests that running speed was unlikely to contribute to the findings. For ANOVAs of the electrophysiology, only rats with correctly placed electrodes (ATN, PFC, and HPC) were included. Repeated-measures polynomial trend analysis was conducted on these data. Hence the specific sample sizes and degrees of freedom for electrophysiological analyses varied, depending on correct electrode location and relevant ipsilateral electrode pairings across the three brain structures (ATN, PFC, and HPC). The LFP analyses focussed on a frequency range that centred on the 8.5 Hz optogenetic stimulation frequency (6.8-10.3 Hz) extracted from the 2-14 Hz frequency band.

## 3. Results

### 3.1 MTT lesions, opsin expression and implants

After three lesion exclusions, and one exclusion due to a misplaced optrode, the rat numbers were: opsin-MTT=12; opsin-Sham=9; and non-opsin MTT=2. Photomicrographs of Luxol blue with cresyl violet stain in a Sham-lesion example and two MTT lesion examples are shown in Fig. 2A. The relevant amount of MTT damage in clinical cases of amnesia is not known and is relatively uncertain in terms of the specificity of the brain damage or the extent of bilateral involvement (e.g. Carlesimo et al., 2007). In our hands, cresyl violet staining alone can give an impression of a complete lesion, but myelin fibre remnants can remain with highly localised lesions. We have found bilateral MTT lesions do not need to be 100% to produce spatial working memory deficits (Perry et al., 2018). Here, to increase the likelihood of severe and persistent deficits during control conditions, we chose the arbitrary inclusion criterion of at least 80% damage to the MTT in each hemisphere based on the Luxol blue staining. For the 12 rats retained in the opsin-MTT group, one rat had 100% MTT lesion on the left and 83% lesion on the right (Fig. 2A, middle panel); all the other 11 rats had 100% bilateral MTT lesions (Fig. 2A, right panel). For the two opsin-MTT rats that were excluded, one had 96% damage on the left but only 63% contralateral damage; the other had 67% and 80% damage on each side respectively. Of the three non-opsin control rats, two had 100% bilateral MTT lesions but the third was excluded due to 74% and 79% damage on the left and right MTT. Placement of electrodes in dorsal hippocampal CA1 and prelimbic cortex, and the optrode in the dorsolateral ATN, are shown in Fig. 2B and 2C. Expression of channelrhodopsin is shown by red fluorescence (mCherry-positive cells) and co-expression with CaMKIIα immunofluorescence indicated viral vector transduction in glutamatergic ATN neurons specifically (Fig. 2B and 2D) and in terminal regions (Fig. 2E: frontal cortex, top left panel; subiculum, bottom left panel; and retrosplenial cortex, top and bottom-right panels). Electrode locations were verified with cresyl violet stain and only data from correctly placed electrodes were retained for analysis.

### 3.2. Spatial working memory after MTT lesions

Prior to optogenetic stimulation, we confirmed that the 12 opsin-MTT rats showed persistent impairment in spatial working memory in the 12-arm maze compared to the 9 opsin-Sham rats (Fig. 2F). This was evident for re-acquisition after lesion surgery (i, Group, F = 5.2, df = 1/19, P = 0.01); after implantation surgery (ii, F = 42.3, df = 1/19, P < 0.001); and the four days before the start of optogenetic TBS (iii, F = 30.1, df = 1/19, P < 0.001; Cohen’s d = 3.318, 95% CI = 1.942-4.657).

### 3.3 Exogenously-triggered optogenetic stimulation and spatial working memory

After demonstrating stable memory impairments in rats with MTT lesions, we next examined the effects of a regular pattern of exogenously-triggered, bilateral optogenetic TBS at 8.5 Hz. Blue-light TBS produced a striking improvement in spatial working memory in the opsin-MTT group, relative to both no stimulation and a regular pattern of orange-light TBS during the first period of optogenetic stimulation (Fig. 3A). For the opsin-MTT group, a large effect size of blue-light stimulation [B] was evident (Cohen’s d = 1.667, 95% CI = 0.762-2.542) when the errors made were averaged over all the sessions with blue-light TBS compared to their performance on sessions with the orange-light [O] TBS (two-tailed paired t-test, t = 5.774, df = 11, P < 0.001). There was no effect of blue light on spatial working memory in the 9 opsin-Sham rats. The effect in the opsin-MTT group was supported by an ANOVA across all conditions, which produced a significant 3-way interaction for Group [MTT vs Sham] x Colour phase [B vs O] x Stimulation [Days with any stimulation vs days with no stimulation], F = 28.7, df = 1/19, P < 0.001. Colour phase refers to the aggregate periods of stimulation and no stimulation for each Colour (i.e., over the first and last four days for the orange phase; and the middle eight days for the blue phase). Higher-order interactions involving the order of test days and all other comparisons within the opsin-MTT group and within the opsin-Sham group were not significant. With respect to the performance of individual rats in the opsin-MTT group, the average reduction in errors with exogenously-triggered blue light TBS (Fig. 3B) reached statistical significance in 10 out of 12 of these rats when assessed by the Reliable Change Index (Jacobson and Truax, 1991).

**Fig. 3.**
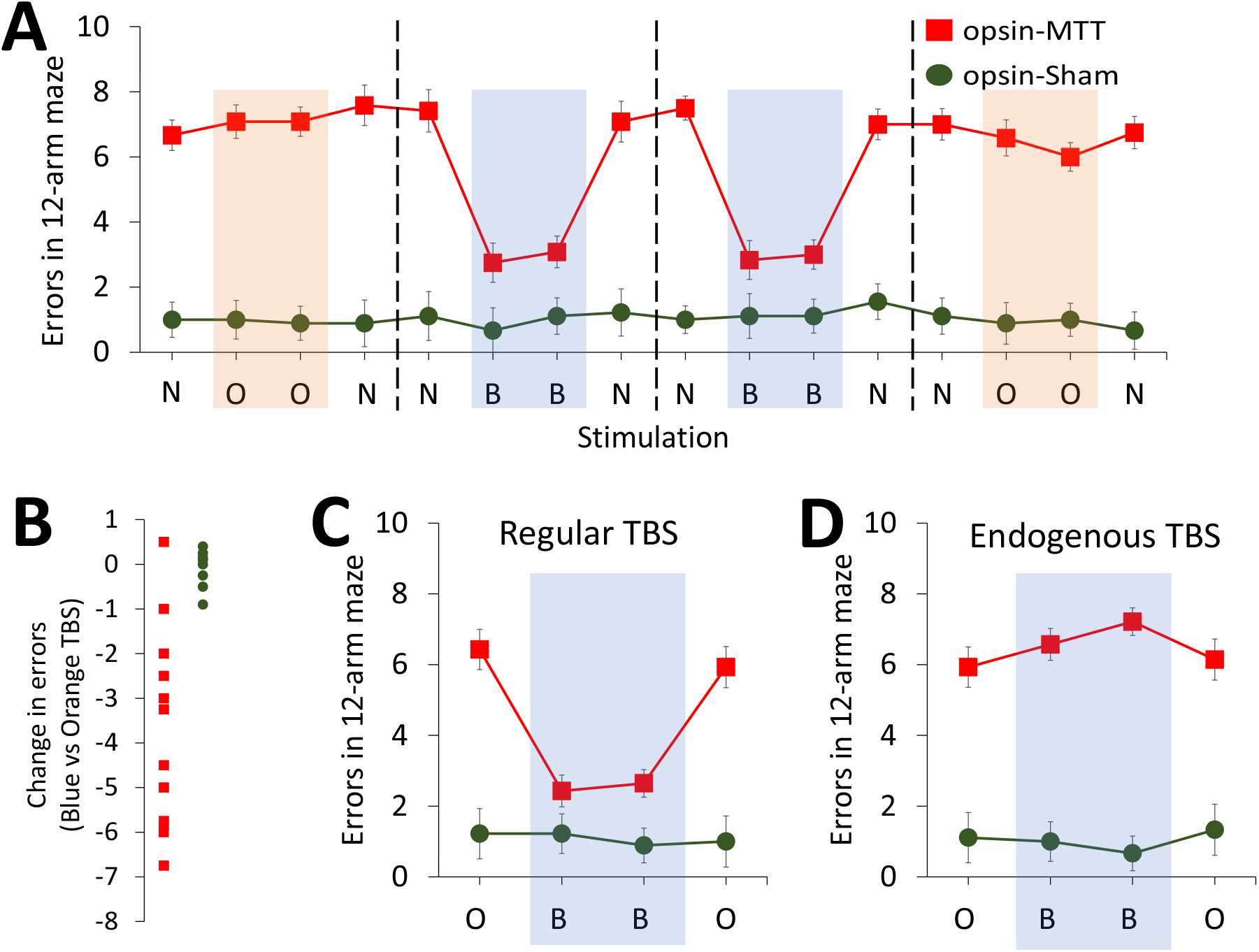
Optogenetic activation of ATN neurons with exogenously-triggered regular blue light theta-burst stimulation (TBS) improves spatial memory in rats with MTT lesions. **A,** Mean spatial working memory errors for the two groups on individual days. Exogenously-triggered regular TBS with blue-light (B) ameliorated the spatial working memory impairment in the opsin-MTT group. **B,** Based on the averaged errors over the four days of each light-stimulation condition shown in A, spatial memory errors were significantly reduced by blue-light stimulation in 10 of 12 opsin-MTT rats (red squares= opsin-MTT; green dots = opsin-Sham). **C,** Improved spatial working memory in the opsin-MTT group with regular blue-light TBS was confirmed in the replication experiment. **D,** By contrast to exogenously-triggered regular blue-light TBS, the opsin-MTT group continued to make substantially more errors than the opsin-Sham group when endogenously-triggered blue-light TBS was used. Error bars = ±SEM.

The beneficial effect on spatial working memory in the 12-arm maze due to exogenously-triggered blue-light optogenetic TBS of the ATN in the opsin-MTT group was replicated when the OBBO sequence was used (Fig. 3C; Group x Colour, F = 36.5, df = 1/19, P < 0.001; B vs O in the opsin-MTT group, Cohen’s d = 2.113, 95% CI = 1.061-3.138).

The two rats with MTT lesions but viral vector infusions that did not include the ChR2(HR) construct (i.e. non-opsin controls) showed unaltered memory performance during exogenous TBS. For example, in the two periods of testing with exogenously-triggered TBS, the mean score on orange days was 8.5 and 6 errors, respectively, and 8 and 7 errors respectively on blue days. So, the improved performance in the opsin-MTT group was unlikely the result of non-specific effects of the viral vector, per se.

### 3.4. Exogenously-triggered optogenetic stimulation and electrophysiology

In the first period of optogenetic stimulation, focusing on the control condition (i.e. days with orange light stimulation; Fig. 4A), MTT lesions compared to Sham lesions reduced ATN PSD (Group, F = 28.8, df = 1/19, P < 0.001), ATN-HPC coherence (F = 5.43, df = 1/17, P = 0.03) and HPC-PFC coherence (F = 9.9, df = 1/11, P = 0.005). No other effects reached significance (all other P-values > 0.10).

**Fig. 4.**
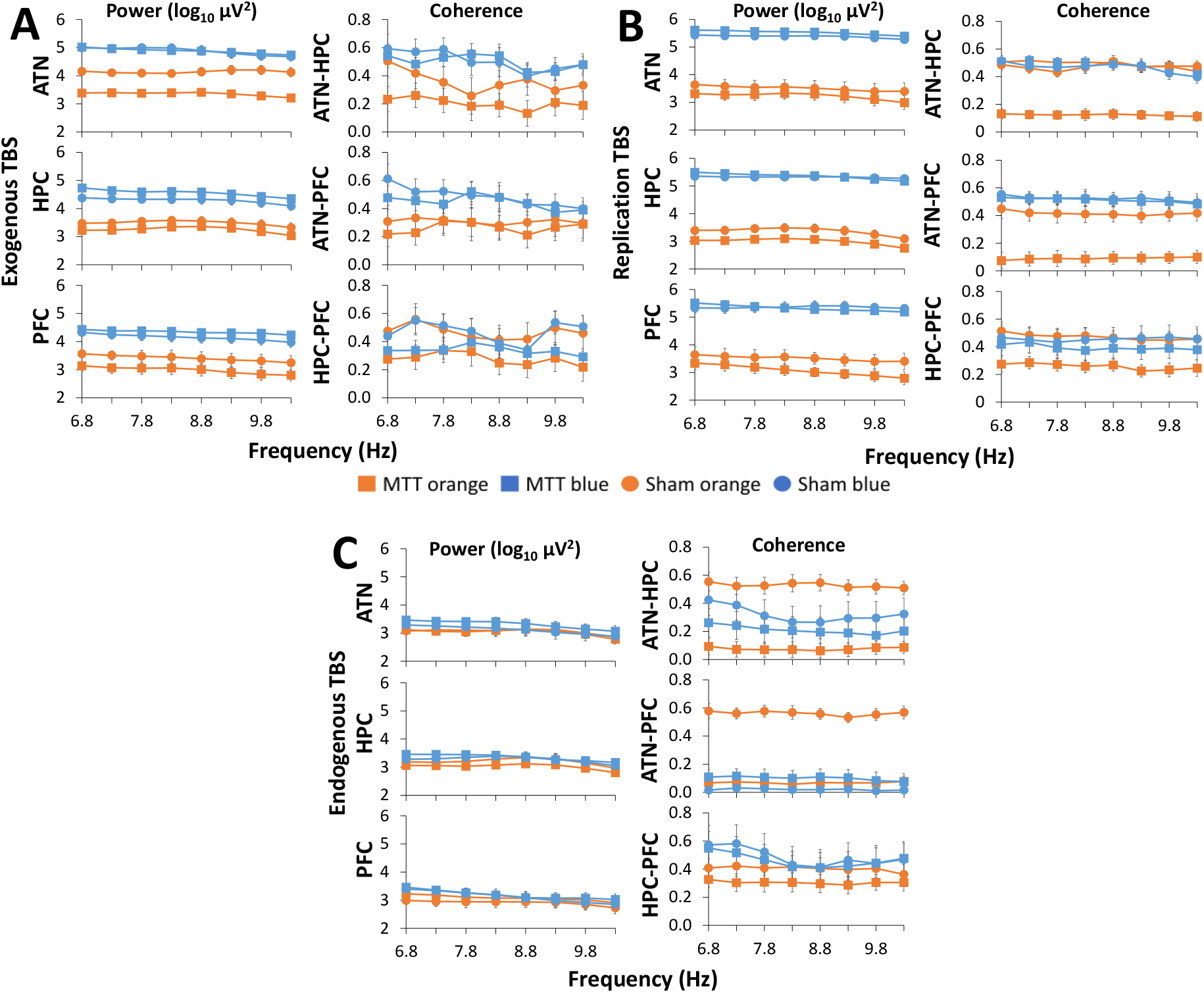
Optogenetic activation of ATN neurons using exogenous, but not endogenous, TBS enhances neural oscillatory activity. Exogenously-triggered optogenetic TBS was applied for the data shown in **A** and **B**. **A,** Power spectral density (PSD) and coherence in the 6.8 Hz to 10.3 Hz range for the aggregated electrophysiology for the last 8 correct choices for B and O days shown in Fig. 3**A**. **B,** PSD and coherence from the replication experiment (last 8 correct choices for B and O days shown in Fig. 3**C**). **C,** Endogenously-triggered blue-light TBS did not alter PSD in the ATN, HPC, or PFC in either group (last 8 correct choices for B and O days shown in Fig. 3**D**); the only effect on coherence was to reduce ATN-PFC coherence in the Sham group. Error bars = ±SEM.

During this first period of TBS, exogenously-triggered regular blue-light TBS increased PSD for both opsin-MTT and opsin-Sham groups in the ATN (Fig. 4A), HPC and PFC compared to orange-light TBS (Colour [i.e., B vs O stimulation days]: ATN, F = 14.8, df = 1/19, P < 0.001; HPC, F = 43.5, df = 1/17, P < 0.001; PFC, F = 21.8, df = 1/11, P = 0.001; df varied with the number of correct electrode placements). In the ATN, the PSD increase was larger in the opsin-MTT group (Group x Colour, F = 6.2, df = 1/19, P = 0.02; opsin-MTT group, B vs O, Cohen’s d = 1.950, 95% CI = 0.953-2.919). The PSD increase was similar in HPC and PFC for both opsin-MTT and opsin-Sham groups (Group x Colour, respectively, F = 2.5, df = 1/17, P = 0.13; and F = 2.2, df = 1/11, P = 0.16).

The blue-light TBS also increased ATN-HPC coherence in both groups (Colour, F = 25.1, df = 1,17, P < 0.001; Group x Colour, F = 2.7, df = 1/11, P = 0.12; Fig. 4A). Blue light stimulation had a similar effect on ATN-PFC coherence (F = 37.0, df = 1/11, P < 0.001), but more so in the opsin-MTT group (Group x Colour, F = 8.7, df = 1/11, P = 0.01;). There was no significant effect of blue light for HPC-PFC coherence (F = 2.1, df = 1/11, P = 0.17).

For the replication with exogenously-triggered TBS, orange light TBS did not reveal PSD differences between the sham and lesion groups in any of the three structures (Group effects, all P > 0.14; Fig. 4B). However, the opsin-MTT group now showed reduced coherence relative to the opsin-Sham group for all three electrode pairs in the orange-light condition (Group: ATN-HPC, F = 47.26, df = 1/19, P < 0.001; ATN-PFC, F = 38.44, df = 1/10, P < 0.001; HPC-PFC, F = 87.03, df = 1/10, P < 0.001).

The exogenously-triggered blue-light TBS during the replication again increased PSD in all three structures (Colour: ATN, F = 634.02, df = 1/19, P < 0.001; HPC, F = 615.48, df = 1/17, P < 0.001; PFC, F = 321.50, df = 1/10, P < 0.001; Fig. 4B). The PSD increase in HPC and PFC was larger in the MTT group (HPC, Group x Colour, F = 5.43, P = 0.03; PFC, Group x Colour, F = 3.31, df = 1/10, P = 0.05). The blue-light TBS produced a large increase in ATN-HPC coherence in the opsin-MTT group, but not the opsin-Sham group (Colour, F = 60.84, df = 1/17, P < 0.001; Group x Colour, F = 53.28, df = 1/17, P < 0.001). There was also a large increase in ATN-PFC coherence in the opsin-MTT group and a smaller increase in the opsin-Sham group (ATN-PFC, Colour, F = 80.49, df = 1/10, P < 0.001; Colour x Group, F = 28.58, df = 1/10, P < 0.001). Regular blue-light TBS also increased HPC-PFC coherence in the opsin-MTT group, but not in the opsin-Sham group (Colour, F = 2.26, df = 1/10, P = 0.16; Group x Colour, F = 3.61, df = 1/10, P < 0.05).

### 3.5 Endogenously-triggered optogenetic stimulation

By contrast to the spatial working memory improvement produced by the regular TBS pattern in the opsin-MTT group, these rats showed no reduction in errors when blue-light stimulation was triggered endogenously from the rat’s own HPC theta (Fig. 3D). That is, the memory impairment in rats with a lesion relative to rats with Sham lesions was not changed (MTT vs Sham Group main effect, F = 80.9, df = 1/19, P < 0.001; Colour [B vs O], F = 0.6, df = 1/19, P = 0.44; Group x Colour, F = 4.1, df = 1/19, P = 0.06). During stimulation with orange light, rats with MTT lesions again showed reduced ATN-HPC (F = 100.56, df = 1/17, P < 0.001) and ATN-PFC coherence (F = 85.77, df = 1/10, P < 0 .001), but there were no lesion effects for either HPC-PFC coherence (F = 3.26, df = 1/11, P > 0.1) or any PSD measures (all P > 0.3). The lack of memory improvement by endogenously triggered blue-light TBS was accompanied by no changes in rhythmic electrophysiology for the opsin-MTT group (Fig. 4C). There was no change in PSD in the ATN, HPC, or PFC in either group (all F < 1.2, all P > 0.3). Endogenously-triggered stimulation had no reliable effects for ATN-HPC or HPC-PFC coherence in either group (Colour, and Group x Colour, all F < 4.88, P > 0.05). While this TBS protocol had no effect on ATN-PFC coherence in the opsin-MTT group, it did substantially reduce ATN-PFC coherence in the opsin-Sham group (Group x Colour, F = 43.0, df = 1/11, P < 0.001).

### 3.6. Impact of regular optogenetic stimulation on Zif268 immediate early gene activation

The neural impact of exogenously-triggered regular optogenetic stimulation of the AV on the expression of Zif268 protein across key structures in the extended hippocampal memory system is shown in Fig. 5. In the hemisphere receiving orange-light TBS (i.e., no opsin activation), the opsin-MTT group showed reduced Zif268 expression in the anterior and posterior granular ‘b’ region of the retrosplenial cortex (RSC; Rgb) and the subiculum (Sub) compared to the opsin-Sham group (Group: Rgb, F = 169.58, df = 1,19, P < 0.001; Sub, F = 6.75, df = 1/19, P = 0.02). Compared to orange-light TBS, blue-light TBS clearly increased Zif268 expression in both opsin-MTT and opsin-Sham groups in structures that are directly innervated by ATN efferents, such as the RSC, Sub and dorsal PFC (Colour: Rgb, F = 169.58, df = 1/19, P < 0.001; Rga, F = 52.1, df = 1/19, P < 0.001; Sub, F = 128.4, df = 1/19, P < 0.001). Blue-light TBS also increased expression in the medial PFC Layer VI of the prelimbic cortex (Colour, F = 17.9, df = 1/19, P < 0.001), and Layer V of all three prefrontal cortex regions examined (Colour, F > 22.1, df = 1/19, P < 0.001). An increase with blue-light stimulation was also found in dorsal hippocampal area CA1 (Colour, F = 69.7, df = 1/19, P < 0.001), which does not receive direct afferents from the ATN. Other regions/layers in the PFC, HPC and RSC were not affected by the optogenetic TBS (Colour, all P > 0.05). No Colour x Group interactions were found (all P > 0.09).

**Fig. 5.**
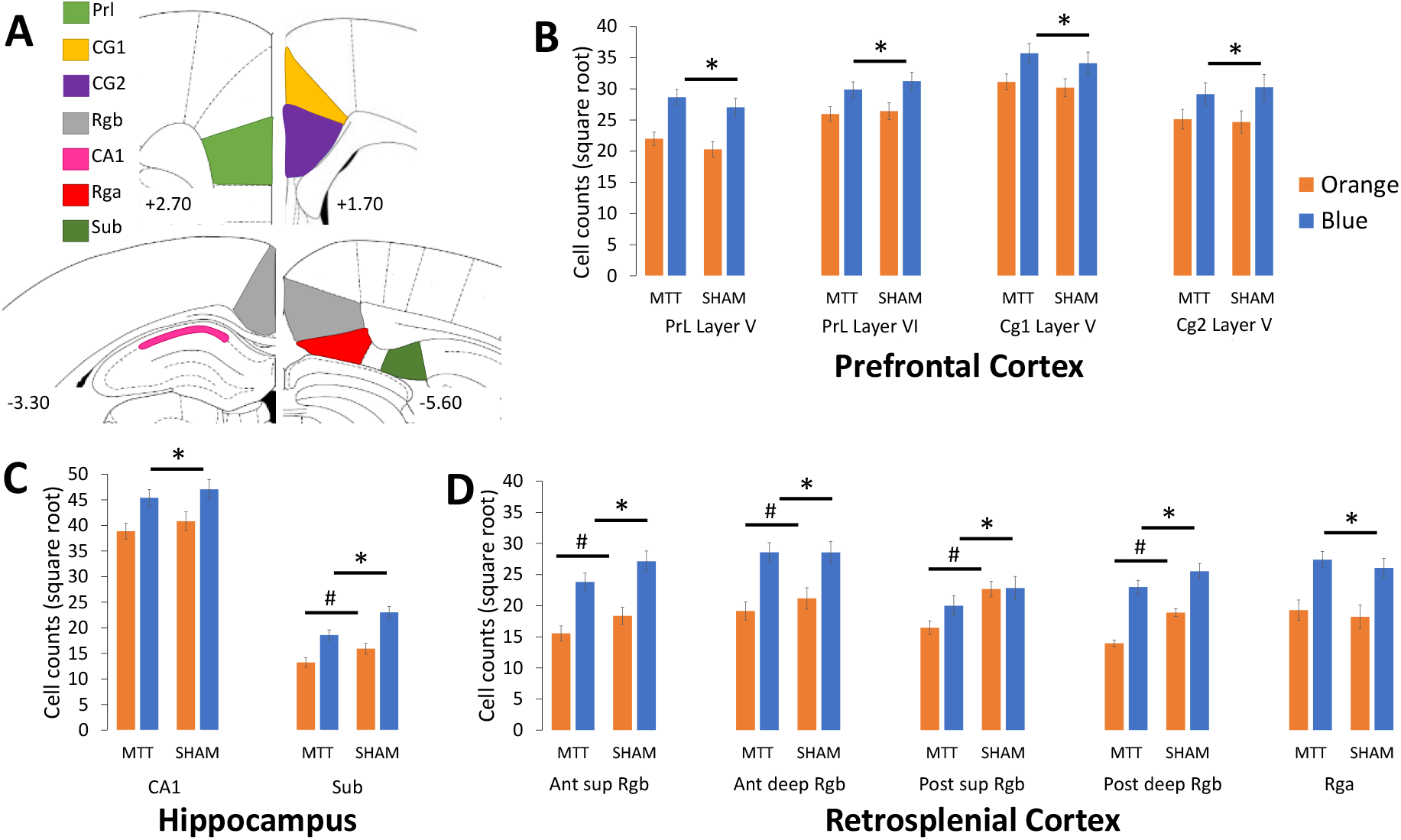
Zif268 expression in memory-related structures is reduced by MTT lesions and increased by exogenously-triggered regular blue-light TBS. **A,** Seven brain regions were assessed after rats foraged for food in an open field when they received exogenously-triggered TBS using blue-light in the ATN in one hemisphere and orange-light in the contralateral ATN. **B,** Expression in the prefrontal cortex. **C,** Expression in CA1 and subiculum. **D**, Expression in the retrosplenial cortex (Ant = anterior; post = posterior; sup = superior layers; deep = deep layers). Other abbreviations: CA1, hippocampal CA1; Cg1, cingulate cortex 1; Cg2, cingulate cortex 2; PrL, prelimbic cortex; Rga, retrosplenial granular a cortex; Rgb, retrosplenial granular b cortex; Sub, dorsal subiculum. Values are relative to bregma. Error bars = ±SEM. # denotes differences between the sham and MTT groups following orange light; * denotes changes related to the blue light condition observed in both groups.

## 4. Discussion

We examined whether selectively stimulating ATN neurons would improve memory performance in rats. The findings reported here demonstrate that optogenetic stimulation focused on AV glutamatergic neurons using an exogenously-triggered regular pattern of TBS at 8.5 Hz acutely improves spatial working memory in rats with MTT lesions. This novel outcome was found despite the rats showing long-term impairment as a consequence of permanent MTT damage. Improved memory was closely linked to the presence of optogenetic stimulation as impaired performance immediately returned on sessions when there was either no stimulation or control light stimulation (i.e. orange light). The improved memory performance in the opsin-MTT group with exogenously-triggered optostimulation was accompanied by broad-band increases in PSD in the ATN, dorsal HPC and PFC, as well as increased ATN-HPC and ATN-PFC coherence. We replicated the acute benefits associated with exogenous optogenetic TBS activation in rats with MTT lesions; this replication adds confidence in the robustness of the observed effects. Relative to control (i.e., orange light) TBS, blue-light TBS also revealed a within-subject increase in Zif268 expression across an impressive array of neural structures that collectively operate as an extended memory system. The increased IEG expression was evident in dorsal hippocampal CA1, dorsal Sub, RSC, and PFC. By contrast, endogenously-triggered TBS based on the rat’s hippocampal theta produced no improvement in spatial memory, which was coupled with a lack of any effects on PSD or rhythmic coherence across the ATN-HPC-PFC axis in the opsin-MTT group.

Vann and colleagues have proposed a model in which the MTT influences distal thalamo-cortical and hippocampal-cortical interactions by optimising memory-related hippocampocortical oscillations (Dillingham et al., 2019, 2021). We provide some support for this model from the effects of MTT lesions on rhythmic activity during spatial working memory testing in a radial arm maze. In the orange-light (i.e., control) condition, rats with MTT lesions showed consistently poorer ATN-HPC rhythmic coherence on all three test occasions, together with poorer HPC-PFC coherence in both the second (replication) period of exogenous stimulation and during the period of endogenous stimulation. These coherence effects suggest that there was no carry-over benefit of prior effective blue-light optostimulation on subsequent rhythmic activity when ineffective exogenously-triggered blue-light or orange-light stimulation was used. The effects of MTT lesions on PSD were limited, however, with only a reduction in the ATN during the first period of exogenous stimulation. Previously, Dillingham and colleagues reported that, during locomotion, MTT lesions disrupted rhythmic activity by attenuating peak theta in the HPC and RSC and producing abnormally increased HPC-RSC coherence (Dillingham et al., 2019). We did not examine HPC-RSC coherence, so we cannot know if this difference (reduced coherence in our study vs increased coherence in their study) depends on the different task demands or the specific memory structures examined. Nonetheless, we provide memory-related electrophysiological evidence that information conveyed by the MTT modulates hippocampal-cortical circuits.

Whereas the impact of MTT lesions was associated with impaired coherence across the ATN-HPC-PFC axis, a broader range of changes, which included increased power across all three structures, was associated with restored spatial memory with exogenous blue-light optostimulation of the ATN. This evidence suggests that the ATN, and perhaps the AV subregion especially, provides the primary mechanism for modulatory control by the MB-ATN axis on distal regions of the extended hippocampal memory system. For the opsin-MTT group, improved spatial working memory with blue-light TBS may be causally related to rhythmic changes in the memory system. Endogenously-triggered blue-light TBS did not change either PSD or coherence across the ATN-HPC-PFC axis and, consistent with this, had no effect on memory in the opsin-MTT group. The beneficial effects of regular exogenously-triggered optostimulation could have resulted from classic short-term potentiation (STP) or frequency potentiation (FP) producing broadband power changes. By contrast, inherently irregular endogenously-triggered stimulation would not produce either STP or FP. STP typically refers to the initial large potentiation observed during the induction of long-term potentiation, but can be evoked in isolation (i.e. without long term potentiation) through the use of weak stimulation (Lisman, 2017). Moreover, it has been suggested that STP may contribute to working memory performance (Erickson et al., 2010; Lisman, 2017). FP is the increase in synaptic responses during low frequency (3-20 Hz) repetitive stimulation. It is thought to depend on the transient increase of neurotransmitters due to residual calcium at the presynaptic terminal (Papatheodoropoulos and Kostopoulos, 2000) and its hippocampal impairment correlates with aged memory dysfunction (Landfield, 1993). Like STP, FP would produce increased transmission in all frequency bands. Note that the effect of this increase in neurotransmitters at the synapse with FP is much longer than a theta cycle, and usually builds steadily over the first few pulses of a train. Whatever the reason behind the differences between regular exogenously-triggered TBS and irregular endogenously-triggered TBS in the current study, the difference between their effects bears some resemblance to an earlier study by McNaughton et al. (2006). They found that regular electrical stimulation using a supramammillary bypass circuit to manipulate hippocampal theta reinstated PSD and theta rhythmicity in the hippocampus and improved spatial memory. This did not occur in their study when irregular stimulation but at the same frequency was used.

The effects of MTT lesions on Zif268 expression were evident in the hemisphere that received orange light (i.e., control) stimulation. MTT lesions reduced expression in the Rgb cortex and the dSub. The reduced IEG expression in the opsin-MTT group was similar to previous work on the distal effects of MTT lesions, although their impact has included more structures in some studies (Vann and Albasser, 2009; Vann, 2013; Frizzati et al., 2016; Perry et al., 2018). It is possible that the unilateral blue-light stimulation may have had some impact in the contralateral hemisphere, but we cannot be certain because we had no comparison without blue-light stimulation. Our semi-quantitative assessment was similar to previous studies on this phenomenon. Future studies could specify these changes in a more quantitative way by using unbiased stereology.

Zif268 immunostaining revealed that exogenously-triggered optogenetic activation of ATN neurons produced marked increases in IEG activation in structures that have been identified as key components in the extended memory network (Aggleton, 2008; Aggleton and Nelson, 2015). Increased expression was found in structures that are directly innervated by ATN efferents, including the anterior and posterior RSC, Sub, and PFC (Bubb et al., 2017; Mathiasen et al., 2017). The assumption that this was due to direct effects of stimulation is supported by observations of ChR2 expression in terminals in these regions. In addition, increased Zif268 expression after blue-light optogenetic TBS was observed in dorsal hippocampal CA1. Unlike the ventral CA1 (de Lima et al., 2016), the dorsal CA1 does not receive direct input from the ATN. So, this suggests that TBS stimulation with blue light was able to alter system-wide functionality and adds further evidence of both direct and indirect influences by the ATN across a distributed network (Bubb et al., 2017; Harland et al., 2014). The finding that optogenetic activation also increased Zif268 expression in the opsin-Sham group in the same structures shows that the effects of optostimulation of ATN neurons are not dependent on a system that has been disrupted by MTT lesions. These findings suggest a broader functional impact of regular 8.5 Hz TBS optostimulation of the ATN on other memory structures than was reported after 130 Hz electrical stimulation of the ATN in intact rats (Hamani et al., 2010).

In the current study, optogenetic stimulation was maintained from the start until the end of the working memory trial in rats that had been trained in the same maze environment on many occasions. The restorative effects of optostimulation in rats with MTT lesions may reflect enhanced acquisition, temporary storage and/or retrieval of spatial information experienced within a given trial or the enhanced processing of stable and well-established long-term representations of the spatial environment. The ATN influence both working and reference spatial memory, whereas the MTT appears to be primarily involved in working memory processes (Perry et al., 2018). Future work could assess these options and whether the beneficial effects of the optogenetically stimulated activity of ATN neurons generalise to other memory tasks associated with the MB-ATN axis, such as memory for temporal order (Mitchell and Dalrymple-Alford, 2005; Dumont and Aggleton, 2013; Wolff et al., 2006; Nelson, 2021). Paired-associate learning tasks provide additional options to consider (e.g., Gilbert and Kesner, 2002; Hunsaker et al., 2006; Gibb et al., 2006). It is possible that the ATN’s strategic position among memory structures is such that it contributes to interactions among three memory systems, that is, an event-based memory system (including spatial working memory), a knowledge-based (consolidated) memory system, and a rule-based (integration) memory system (Hunsaker and Kesner, 2018). The current findings support the prospect of additional studies on the optogenetic activation of ATN neurons and to address the possibility that the ATN’s role in memory extends beyond the classic areas of spatial memory and memory for temporal order (Nelson, 2021). Unlike lesion or suppression, evidence from the direct activation of the ATN may provide a more complete account of the role of this region in memory.

The placement of the optic fibres was focused on the AV subnucleus in the current study. It is possible that the observed optostimulation effects derived primarily from an influence on neurons in this region. There is evidence to support the idea of three parallel ATN circuits across the extended memory system, broadly defined by the three ATN subnuclei (Aggleton et al., 2010). If our findings were primarily due to optostimulation of the AV, then this reinforces the proposal that the AV, in particular, plays an important role in supporting spatial navigation. We are unable, however, to specify that there was no contribution from AM neurons or even nearby non-ATN neurons. While the AM is considered as being an important site of convergence that sends HPC-diencephalic information to frontal areas, it is also involved in the propagation of theta signals in the Papez circuit (Aggleton et al, 2010; Jankowski et al., 2013). Understanding the unique and overlapping ways that the AV and AM contribute to memory processing will require more selective experimental manipulations.

Our study provides compelling evidence for an active role of the ATN in a distributed memory network. Previously, electrical stimulation has been used, but this manipulation is less specific and resulted in mixed effects in rats (Hamani et al., 2010, 2011), although there is recent evidence that electrical stimulation of the ATN improves working memory in drug-resistant epilepsy patients (Liu et al., 2021). Here, we showed that optogenetic activation of ATN glutamatergic projection neurons using exogenously-triggered regular TBS immediately improved spatial working memory in rats with MTT lesions that otherwise showed profound and persistent deficits. This optogenetic stimulation also improved key elements of rhythmic electrical activity and increased IEG activation across memory-related structures. That is, the independent activity of ATN projection neurons significantly improved function in the network of brain structures that support spatial memory. The lack of impact on memory or rhythmic activity when optogenetic stimulation was endogenously-triggered by hippocampal theta suggests that the ATN’s involvement in spatial memory is not primarily due to their hippocampal inputs.

Taken together, our study provides direct evidence that the ATN provide far more than simple relays of hippocampal information. Such evidence contributes an additional example of thalamic control of cognition (Halassa and Kastner, 2017; Wolff and Vann, 2019). While much more work is needed, this evidence suggests that gene therapy focused on a relatively small brain structure, the ATN, may be a feasible target for memory rehabilitation. It may counter amnesia caused by a range of brain insults that influence ATN integrity, from early life hypoxia-ischaemia, adult thalamic injury, to Alzheimer’s neuropathology (Aggleton, 2014; Barnett et al., 2018).

## Acknowledgements

JCD-A (PI), LCP-B (co-PI), SMH and NMcN gratefully acknowledge project funding, as well as viral vector support, and PhD scholarship (awarded to SCB) from Brain Research New Zealand - Rangahau Roro Aotearoa, a national Centre of Research Excellence. JCD-A, BALP and SCB acknowledge additional support from the University of Canterbury, New Zealand.

## Author contributions

JCD-A, concept, project design, funding, analyses and primary drafts. SCB, project design, conducting surgery and behavioural experiments, histology, analyses, and primary drafts. LP-B, project design, funding. BP, concept, funding. CKY, analyses. HW, viral vector materials. SH, project design, funding. NMcN, project design, analyses, and primary drafts. All authors interpreted data and edited the manuscript.

## References

Aggleton, J. P., Brown, M. W., 1999. Episodic memory, amnesia, and the hippocampal–anterior thalamic axis. Behavioral and Brain Sciences, 22(3), 425–444.

Aggleton, J. P., 2008. Understanding anterograde amnesia: disconnections and hidden lesions. The Quarterly Journal of Experimental Psychology, 61(10), 1441–1471.

Aggleton, J. P., O’Mara, S. M., Vann, S. D., Wright, N. F., Tsanov, M., Erichsen, J. T., 2010. Hippocampal-anterior thalamic pathways for memory: uncovering a network of direct and indirect actions. European Journal of Neuroscience. 31, 2292–2307.

Aggleton, J. P., 2014. Looking beyond the hippocampus: old and new neurological targets for understanding memory disorders. Proceedings of the Royal Society B: Biological Sciences, 281(1786), 20140565.

Aggleton, J. P., Nelson, A. J., 2015. Why do lesions in the rodent anterior thalamic nuclei cause such severe spatial deficits? Neuroscience and Biobehavioral Reviews, 54, 131–144.

Aggleton, J. P., Pralus, A., Nelson, A. J., Hornberger, M., 2016. Thalamic pathology and memory loss in early Alzheimer’s disease: moving the focus from the medial temporal lobe to Papez circuit. Brain, 139(7), 1877–1890.

Barnett, S. C., Perry, B. A. L., Dalrymple-Alford, J. C., Parr-Brownlie, L. C., 2018. Optogenetic stimulation: Understanding memory and treating deficits. Hippocampus, 28(7), 457–470.

Bubb, E. J., Kinnavane, L., Aggleton, J. P., 2017. Hippocampal–diencephalic–cingulate networks for memory and emotion: An anatomical guide. Brain and Neuroscience Advances, 1, 2398212817723443.

Caulo, M., Van Hecke, J., Toma, L., Ferretti, A., Tartaro, A., Colosimo, C., Romani, G. L., Uncini, A., 2005. Functional MRI study of diencephalic amnesia in Wernicke–Korsakoff syndrome. Brain, 128(7), 1584–1594.

Carlesimo, G. A., Lombardi, M. G., Caltagirone, C., 2011. Vascular thalamic amnesia: a reappraisal. Neuropsychologia, 49(5), 777–789.

Carlesimo, G. A., Serra, L., Fadda, L., Cherubini, A., Bozzali, M., Caltagirone, C., 2007. Bilateral damage to the mammillo-thalamic tract impairs recollection but not familiarity in the recognition process: a single case investigation. Neuropsychologia, 45(11), 2467–2479.

de Lima, M. A. X., Baldo, M. V. C., Canteras, N. S., 2016. A role for the anteromedial thalamic nucleus in the acquisition of contextual fear memory to predatory threats. Brain Structure and Function. 222 (1), 113e129.

Dillingham, C. M., Frizzati, A., Nelson, A. J., Vann, S. D., 2015. How do mammillary body inputs contribute to anterior thalamic function? Neuroscience and Biobehavioral Reviews, 54, 108–119.

Dillingham, C. M., Milczarek, M. M., Perry, J. C., Frost, B. E., Parker, G. D., Assaf, Y., Sengpiel, F., O’Mara, S. M., Vann, S. D., 2019. Mammillothalamic disconnection alters hippocampocortical oscillatory activity and microstructure: implications for diencephalic amnesia. Journal of Neuroscience, 39(34), 6696–6713.

Dillingham, C. M., Milczarek, M. M., Perry, J. C., Vann, S. D., 2021. Time to put the mammillothalamic pathway into context. Neuroscience and Biobehavioral Reviews, 121, 60–74.

Dumont, J. R., Amin, E., Poirier, G. L., Albasser, M. M., Aggleton, J. P., 2012. Anterior thalamic nuclei lesions in rats disrupt markers of neural plasticity in distal limbic brain regions. Neuroscience, 224, 81–101.

Dumont, J. R., Aggleton, J. P., 2013. Dissociation of recognition and recency memory judgments after anterior thalamic nuclei lesions in rats. Behavioral Neuroscience, 127(3), 415.

Erickson, M. A., Maramara, L. A., Lisman, J., 2010. A single brief burst induces GluR1-dependent associative short-term potentiation: a potential mechanism for short-term memory. Journal of Cognitive Neuroscience, 22(11), 2530–2540.

Farina, F. R., Commins, S., 2016. Differential expression of immediate early genes Zif268 and c-Fos in the hippocampus and prefrontal cortex following spatial learning and glutamate receptor antagonism. Behavioural Brain Research, 307, 194–198.

Ferguson, M. A., Lim, C., Cooke, D., Darby, R. R., Wu, O., Rost, N. S., Corbetta, M., Grafman, J., Fox, M. D., 2019. A human memory circuit derived from brain lesions causing amnesia. Nature Communications, 10(1), 1–9.

Frizzati, A., Milczarek, M. M., Sengpiel, F., Thomas, K. L., Dillingham, C. M., Vann, S. D., 2016. Comparable reduction in Zif268 levels and cytochrome oxidase activity in the retrosplenial cortex following mammillothalamic tract lesions. Neuroscience, 330, 39–49.

Gallo, F. T., Katche, C., Morici, J. F., Medina, J. H., Weisstaub, N. V., 2018. Immediate early genes, memory and psychiatric disorders: focus on c-Fos, Egr1 and Arc. Frontiers in Behavioral Neuroscience, 12, 79.

Gilbert, P. E., Kesner, R. P., 2002. Role of rodent hippocampus in paired-associate learning involving associations between a stimulus and a spatial location. Behavioral Neuroscience, 116(1), 63.

Halassa, M. M., Kastner, S., 2017. Thalamic functions in distributed cognitive control. Nature Neuroscience, 20(12), 1669–1679.

Hamani, C., Dubiela, F. P., Soares, J. C., Shin, D., Bittencourt, S., Covolan, L., Carlen, P. L., Laxton, A. W., Hodaie, M., Stone, S. S., Ha, Y., 2010. Anterior thalamus deep brain stimulation at high current impairs memory in rats. Experimental Neurology, 225(1), 154–162.

Hamani, C., Stone, S. S., Garten, A., Lozano, A. M., Winocur, G., 2011. Memory rescue and enhanced neurogenesis following electrical stimulation of the anterior thalamus in rats treated with corticosterone. Experimental Neurology, 232(1), 100–104.

Harding, A., Halliday, G., Caine, D., Kril, J., 2000. Degeneration of anterior thalamic nuclei differentiates alcoholics with amnesia. Brain, 123(1), 141–154.

Harland, B. C., Collings, D. A., McNaughton, N., Abraham, W. C., Dalrymple-Alford, J. C., 2014. Anterior thalamic lesions reduce spine density in both hippocampal CA1 and retrosplenial cortex, but enrichment rescues CA1 spines only. Hippocampus, 24(10), 1232–47.

Hunsaker, M. R., Thorup, J. A., Welch, T., Kesner, R. P., 2006. The role of CA3 and CA1 in the acquisition of an object-trace-place paired-associate task. Behavioral Neuroscience, 120(6), 1252.

Hunsaker, M. R., Kesner, R. P., 2018. Unfolding the cognitive map: The role of hippocampal and extra-hippocampal substrates based on a systems analysis of spatial processing. Neurobiology of Learning and Memory, 147, 90–119.

Jones, M. W., Errington, M. L., French, P. J., Fine, A., Bliss, T. V., Garel, S., Charnay, P., Bozon, B., Laroche, S. and Davis, S., 2001. A requirement for the immediate early gene Zif268 in the expression of late LTP and long-term memories. Nature Neuroscience, 4(3), 289–296.

Kim, E., Ku, J., Namkoong, K., Lee, W., Lee, K. S., Park, J. Y., Lee, S. Y., Kim, J. J., Kim, S. I., Jung, Y. C., 2009. Mammillothalamic functional connectivity and memory function in Wernicke’s encephalopathy. Brain, 132(2), 369–376.

Landfield, P. W., 1993. Impaired frequency potentiation as a basis for aging-dependent memory impairment: The role of excess calcium influx. Neuroscience Research Communications, 13 Suppl. 1, S19–S22.

Lisman, J., 2017. Glutamatergic synapses are structurally and biochemically complex because of multiple plasticity processes: long-term potentiation, long-term depression, short-term potentiation and scaling. Philosophical Transactions of the Royal Society B: Biological Sciences, 372(1715), 20160260.

Liu, J., Yu, T., Wu J., Pan, Y., Tan, Z., Liu, R., Wang. X., Ren, L., Wang, L., 2021. Anterior thalamic stimulation improves working memory precision judgments, Brain Stimulation, doi: https://doi.org/10.1016/j.brs.2021.07.006.

Loukavenko, E. A., Wolff, M., Poirier, G. L., Dalrymple-Alford, J. C., 2016. Impaired spatial working memory after anterior thalamic lesions: Recovery with cerebrolysin and enrichment. Brain Structure and Function, 221(4), 1955–1970.

Mathiasen, M. L., Dillingham, C. M., Kinnavane, L., Powell, A. L., Aggleton, J. P., 2017. Asymmetric cross-hemispheric connections link the rat anterior thalamic nuclei with the cortex and hippocampal formation. Neuroscience, 349, 128–143.

McNaughton, N., Ruan, M., Woodnorth, M. A., 2006. Restoring theta-like rhythmicity in rats restores initial learning in the Morris water maze. Hippocampus, 16(12), 1102–1110.

Mitchell, A. S., Dalrymple-Alford, J. C., 2006. Lateral and anterior thalamic lesions impair independent memory systems. Learning and Memory, 13(3), 388–396.

Nelson, A. J., Vann, S. D., 2017. The importance of mammillary body efferents for recency memory: towards a better understanding of diencephalic amnesia. Brain Structure and Function, 222(5), 2143–2156.

Nelson, A. J., Perry, J. C., Vann, S. D., 2018. The Papez Circuit and Recognition Memory: Contributions of the Medial Diencephalon and Retrosplenial Cortex to What, Where and When Aspects of Object Recognition Memory. In Handbook of Behavioral Neuroscience (Vol. 27, pp. 217–226). Elsevier.

Nelson, A. J., Kinnavane, L., Amin, E., O’Mara, S. M., Aggleton, J. P., 2020. Deconstructing the direct reciprocal hippocampal-anterior thalamic pathways for spatial learning. Journal of Neuroscience, 40(36), 6978–6990.

Nelson, A. J., 2021. The anterior thalamic nuclei and cognition: a role beyond space? Neuroscience & Biobehavioral Reviews, 126, 1–11.

Papatheodoropoulos, C., Kostopoulos, G., 2000. Decreased ability of rat temporal hippocampal CA1 region to produce long-term potentiation. Neuroscience Letters, 279(3), 177–180.

Parr-Brownlie, L. C., Bosch-Bouju, C., Schoderboeck, L., Sizemore, R., Abraham, W., Hughes, S. M., 2015. Lentiviral vectors as tools to understand central nervous system biology in mammalian model organisms. Frontiers in Molecular Neuroscience, 8, 14.

Penke, Z., Morice, E., Veyrac, A., Gros, A., Chagneau, C., LeBlanc, P., Samson, N., Baumgärtel, K., Mansuy, I. M., Davis, S., Laroche, S., 2014. Zif268/Egr1 gain of function facilitates hippocampal synaptic plasticity and long-term spatial recognition memory. Philosophical Transactions of the Royal Society B: Biological Sciences, 369(1633), p.20130159.

Pergola, G., Ranft, A., Mathias, K., Suchan, B., 2013. The role of the thalamic nuclei in recognition memory accompanied by recall during encoding and retrieval: an fMRI study. Neuroimage, 74, 195–208.

Perry, B. A., Mercer, S. A., Barnett, S. C., Lee, J., Dalrymple-Alford, J. C., 2018. Anterior thalamic nuclei lesions have a greater impact than mammillothalamic tract lesions on the extended hippocampal system. Hippocampus, 28(2), 121–135.

Reed, L. J., Lasserson, D., Marsden, P., Stanhope, N., Stevens, T., Bello, F., Kingsley, D., Colchester, A., Kopelman, M. D., 2003. FDG-PET findings in the Wernicke-Korsakoff syndrome. Cortex, 39(4-5), 1027–1045.

Sweeney-Reed, C. M., Buentjen, L., Voges, J., Schmitt, F. C., Zaehle, T., Kam, J. W., Kaufmann, J., Heinze, H. J., Hinrichs, H., Knight, R. T., Rugg, M. D., 2021. The role of the anterior nuclei of the thalamus in human memory processing: Contribution to special issue in Neuroscience & Biobehavioral Reviews, 126, 146–158.

Vann, S. D., 2013. Dismantling the Papez circuit for memory in rats. Elife, 2, e00736.

Vann, S. D., Albasser, M. M., 2009. Hippocampal, retrosplenial, and prefrontal hypoactivity in a model of diencephalic amnesia: Evidence towards an interdependent subcortical-cortical memory network. Hippocampus, 19(11), 1090–1102.

Wolff, M., Gibb, S. J., Dalrymple-Alford, J. C., 2006. Beyond spatial memory: the anterior thalamus and memory for the temporal order of a sequence of odor cues. Journal of Neuroscience, 26(11), 2907–2913.

Wolff, M., Vann, S. D., 2019. The cognitive thalamus as a gateway to mental representations. Journal of Neuroscience, 39(1), 3–14.

Żakowski, W., 2017. Neurochemistry of the anterior thalamic nuclei. Molecular Neurobiology, 54(7), 5248–5263.

Żakowski, W., Zawistowski, P., Braszka, Ł., Jurkowlaniec, E., 2017. The effect of pharmacological inactivation of the mammillary body and anterior thalamic nuclei on hippocampal theta rhythm in urethane-anesthetized rats. Neuroscience, 362, 196–205.

Zotev, V., Misaki, M., Phillips R., Wong C. K., Bodurka, J., 2018. Real-time fMRI neurofeedback of the mediodorsal and anterior thalamus enhances correlation between thalamic BOLD activity and alpha EEG rhythm. Human Brain Mapping, 39, 1024–1042.

